# Remembrance with gazes passed: Eye movements precede continuous recall of episodic details of real-life events

**DOI:** 10.1101/2024.02.29.582757

**Authors:** Ryan M. Barker, Michael J. Armson, Nicholas B. Diamond, Zhong-Xu Liu, Yushu Wang, Jennifer D. Ryan, Brian Levine

## Abstract

Autobiographical memory entails reconstructing the visual features of past events. Eye movements are associated with vivid autobiographical recollection, but this research has yet to capitalize on the high temporal resolution of eye-tracking data. We aligned eye movement data with participants’ simultaneous free recall of a verified real-life event, allowing us to assess the temporal correspondence of saccades to production of episodic and non-episodic narrative content at the millisecond level. Eye movements reliably predicted subsequent episodic – but not non-episodic – details by 250-1100 ms, suggesting that they facilitate episodic recollection by reinstating spatiotemporal context during vivid recollection. Assessing the relationship of oculomotor responses to naturalistic memory informs theory as well as diagnosis and treatment of conditions involving pathological recollection, such as Alzheimer’s disease and post-traumatic stress disorder (PTSD).

## Introduction

Episodic recollection involves re-experiencing the multimodal context of a previous event, including spatiotemporal, emotional, and sensory details (1) with vision as the dominant sensory modality (2). Although altered recollection is a hallmark sign of medial temporal lobe pathology (as in Alzheimer’s disease) and psychopathology (as in intrusive imagery or flashbacks in post-traumatic stress disorder; PTSD), it is difficult to quantify due to its inherently subjective state. Consequently, researchers have leveraged the well-characterized anatomy and physiology of the visual system to assess correlates of conscious experience in recollection (3), allowing for inferences to be made regarding the quality and content of visual memories. Other researchers have found that eye movements carry information about the nature of visual processing that would not be evident from activity in visual sensory regions at encoding and retrieval (4), even in the absence of visual stimuli (5, 6).

A growing body of research has supported a role for unconstrained eye movements in naturalistic autobiographical recollection (7). Compared to laboratory paradigms, these naturalistic paradigms are more closely aligned with real-life normal or disordered function. On the other hand, the use of self-selected lifespan events sacrifices experimental control of memory content, remoteness, and prior rehearsal. Using a staged event (a museum-like tour encoded one week prior to testing; 8), we found that the rate of eye movements was specifically related to the quantified richness and specificity of event memory (9), particularly for individuals who report visually rich autobiographical recollections in general (10).

This research suggests that eye movements are related to recall of real-life episodic details over and above non-episodic details or other cognitive operations involved in complex autobiographical recall. These analyses, however, average over epochs of narrative recall, so they cannot speak to the directionality and temporality of the relationship between eye movements and detail production. The high temporal resolution of eye tracking data, which approximates that of neural signals (11), carries additional information about the timing of cognitive operations. If eye movements actively facilitate visual imagery, setting the stage for recall in time and place (6), they should reliably precede episodic detail production. Alternatively, if eye movements reflect a response to vocalized detail production or non-specific retrieval-related processes, they should be more evident after episodic detail production, or they should show no reliable relationship to the timing of specific narrative content categories.

To address these questions, we aligned the time-series of oculomotor responses to verbal free recall of a controlled real-life event (10) at the millisecond level. Transcribed narrative content was categorized into episodic (internal) or non-episodic (external) details using a reliable coding method (9), allowing us to assess the relationship between eye movements and the episodic vs. non-episodic recall across a sliding window surrounding each detail (i.e., +/- 1500 ms before and after detail onset, or 3000 ms, segmented into 10 time bins). Understanding these sequential relationships would inform theories concerning naturalistic episodic memory mechanisms as well as assessment and treatment of clinical conditions involving episodic recollection.

## Results

Eye movements were amplified in the epochs preceding internal, but not external details. Using a logistic mixed-effects model with a random intercept and slope, we found a significant interaction between saccades and time bin (*χ*^2^(9) = 18.63, *p* = .03). Helmert-coded beta coefficients from the model revealed internal details were preceded by increased eye movements in the second and third bins centered on 625 and 875 ms prior to detail onset (range: -1250 – - 250 ms; *b*’s = 0.09 and 0.10, *SE’*s = 0.04, *Z*’s = 2.25 and 2.53, *p’*s = .02 and .01, *odds ratios =* 1.10 and 1.11 for the second and third bins, respectively) than the average of all subsequent time bins (Figure 1). Indeed, eye movements made after internal detail recall were attenuated, relative to the average.

**Figure 1.**
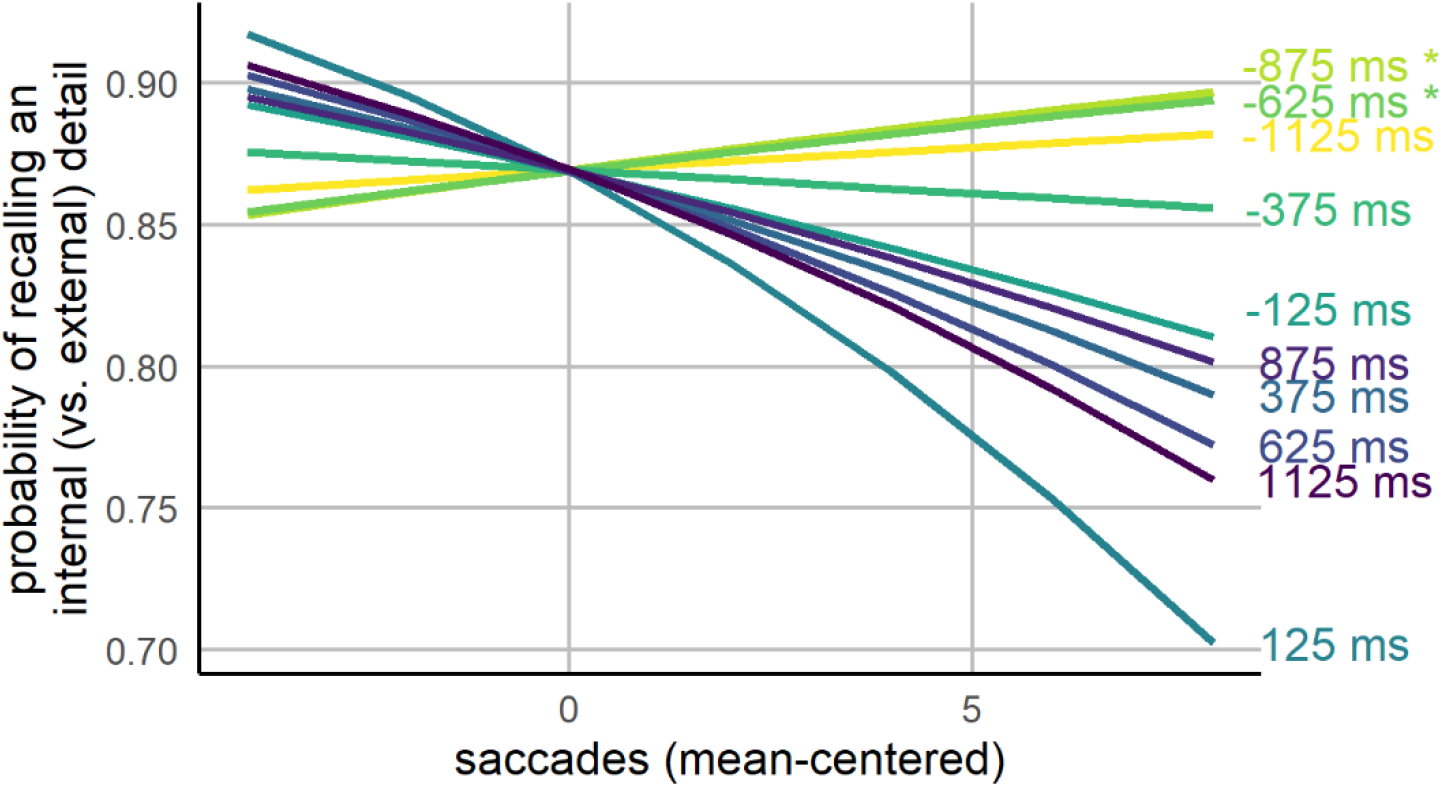
Predicted Detail Type by Timing of Eye Movements *Note*. Probability of an internal (vs. external) detail following saccade plotted as a function of saccade timing. Slope labels indicate midpoint of each time bin. Time bins are colour-coded as described in Figure 3 (see methods). Saccades occurring in the early time bins (i.e., centered on 875 and 625 ms prior to detail onset) predicted the probability of an internal (vs. external) detail; internal details were followed by an attenuation of eye movements.

The nature of the distribution of eye movements in relation to detail production was tested with mixed-effects (random intercept and slope) cubic polynomial regressions on internal and external detail data subsets. The model comprising internal detail data revealed a significant cubic term for time bin (*F*(3, 242.53) = 17.06, *p* < .001), which was a better fit than a simpler, quadratic model (AIC_quadratic_ = 104204, AIC_cubic_ = 104169, BIC_quadratic_ = 104263, BIC_cubic_ = 104236; *χ*^2^(1) = 37.64, *p* = < .001; see Figure 2. The model considering external details did not demonstrate a reliable cubic trend (*F*(3, 155.84) = 1.5459, *p* = .20); model comparison found a linear relationship (*F*(1, 53.94) = 0.03, *p* = .86) best fit the external detail data across time bins as the cubic term did not improve model fit compared to a quadratic model (AIC_quadratic_ = 22443, AIC_cubic_ = 22442, BIC_quadratic_ = 22491, BIC_cubic_ = 22497; *χ*^2^(1) = 2.94, *p* = .08) nor did the quadratic model improve the model’s fit above a linear model (AIC_linear_ = 22442, BIC_linear_ = 22484; *χ*^2^(1) = 1.66, *p* = .20). Finally, a model including eye movements made for both internal and external details confirmed the dissociation in eye movement patterns with a reliable interaction between detail type and a cubic term for time bin (*F*(3, 39050) = 4.24, *p* = .005).

**Figure 2.**
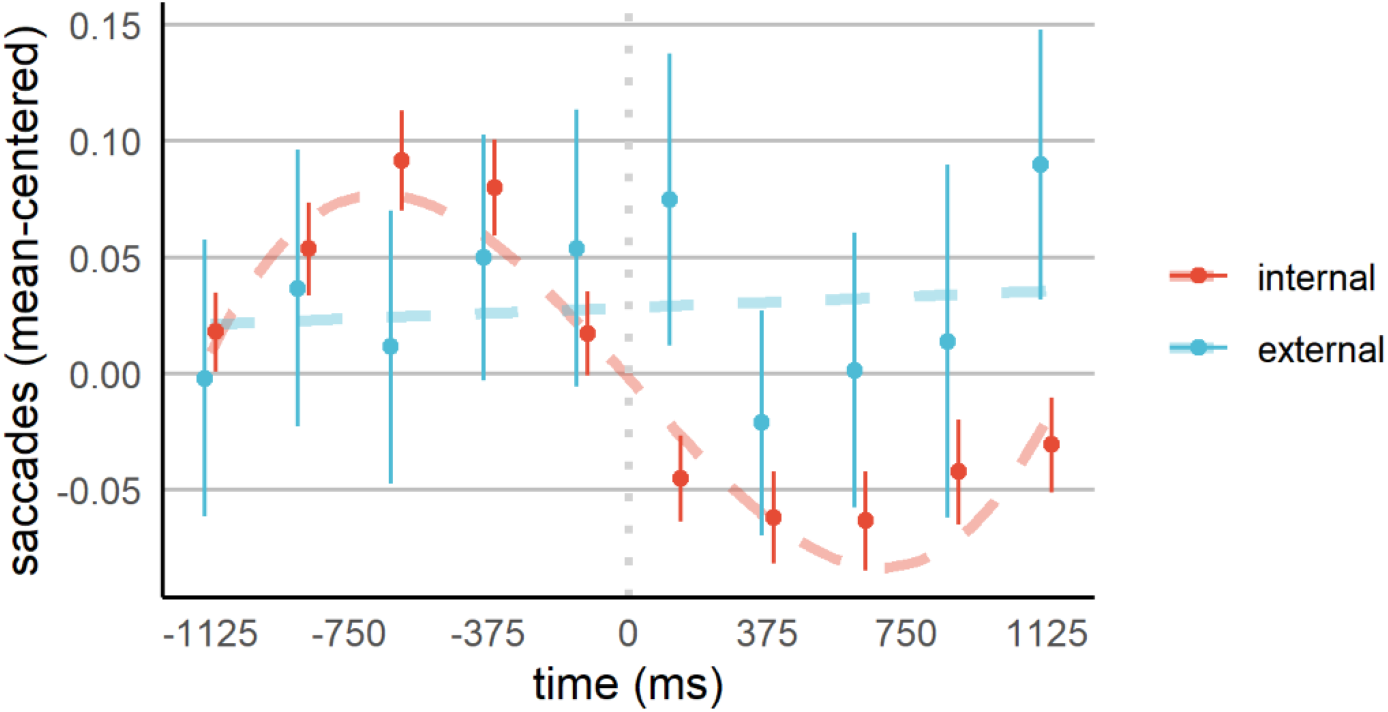
Polynomial and Linear Model Fit to Eye Movements during Internal and External Detail Recall *Note*. Average saccades by time bin are plotted alongside regression model predictions found to best fit data comprising internal details (cubic trend) and external details (linear trend). Data points are plotted at the midpoint of the respective time window. Dashed lines depict model trends. Dotted line represents detail onset. Error bars represent standard error.

## Discussion

We investigated the directional relationship between eye movements and episodic autobiographical memory for real-life events at a fine temporal scale. As predicted, eye movements reliably preceded the recall of episodic autobiographical details. The observed patterns were specific to episodic vs non-episodic content produced within a single narrative protocol. These findings suggest that eye movements are causally—and specifically—related to the production of episodic details in autobiographical free recall.

Our staged event was optimized for engagement of episodic autobiographical memory with respect to novelty, recency, and extensive temporal, spatial, and enacted components that distinguish real-life events from laboratory memory paradigms. The fidelity of staged event recall was demonstrated through high accuracy of the encoded details (8). Given the recency of the event, subjective re-experiencing was vivid yet not excessively rehearsed as compared to self-selected personal events, as is typical in autobiographical memory research.

The observed temporal lag of 1250-250 ms between eye movements and episodic detail production is consistent with prior research in which eye movement indices of memory precede verbal responses (12), allowing for 100-200 ms in speech preparation time (13). These movements may therefore reflect an obligatory retrieval process, such as pattern completion, to reinstate the encoding-related spatiotemporal context necessary for recall of real-life events (6).

The post-retrieval diminishment in eye movements may signal the end of a retrieval cycle whereby the interrogation of the mental representation is no longer required as pre-speech verbal processing ensues. Alternatively, it may reflect the rapid suppression of retrieval to reduce interference that may otherwise weaken the activated representation (5). In this manner, the cycle of saccades and eye fixations promotes the continuous recall of new details.

Free recall of episodic autobiographical details is strongly associated with the functioning of the hippocampus and surrounding medial temporal lobe structures and their connections to cortical regions (14, 15). Our findings suggest an antecedent role of eye movements in a temporal arc that induces visual sensory re-experiencing and autobiographical recollection. Imaging methods with high temporal resolution could be used to test hypotheses concerning the neural sequences of motor, mnemonic, and sensory signals across the autobiographical memory network.

Given a known, sequential relationship between eye movements and everyday verbal memory behavior, the decoupling of such responses could be used as a marker for the deficits in recollection that accompany medial temporal lobe damage, as in Alzheimer’s disease. Conversely, post-traumatic stress disorder (PTSD) entails highly vivid and intrusive recollection of traumatic events as well as functional and structural alterations in visual cortical networks (16). Effective interventions for PTSD entail contextualizing traumatic events and reducing the emotional impact of intrusive visual memories (17). Fine-grain behavioral analysis as reported here could be used to test and refine such interventions.

## Methods

Ninety-one healthy young adult participants (66 females, mean age = 24.77, mean education = 16.56 years) completed an audio-guided museum-style tour of artworks and installations throughout Baycrest Hospital (available at https://osf.io/j25y4/). Verbal free recall of the tour was assessed one week later with simultaneous eye tracking (EyeLink II system, SR Research Ltd; Mississauga, Ontario, Canada) while viewing a blank computer screen.

Recall responses were transcribed via Google Speech-to-Text in Python (https://cloud.google.com/speech-to-text) such the placement of each word on the time series could be determined at the millisecond level. This transcription was annotated with narrative detail categorization as reliably determined by the Autobiographical Interview scoring procedure (12), which categorizes details as internal (i.e., information directly relating to or contextually embedded in the event recollected) or external (i.e., semantic information or information not specific to the event recollected). Conversion of the eye movement data to a series of fixation and saccade events that were time-locked to memory recall (via an auditory tone) was achieved and interrogated with Data Viewer (SR Research Ltd.). Fixations and saccades were defined via EyeLink’s online parser. A 3000 ms sliding window was constructed around each detail onset (see Figure 3). Saccades were summed within ten 750 ms-wide bins, with each bin separated by a lag of 250 ms. Additional methodological details are included in the supplementary materials.

**Figure 3.**
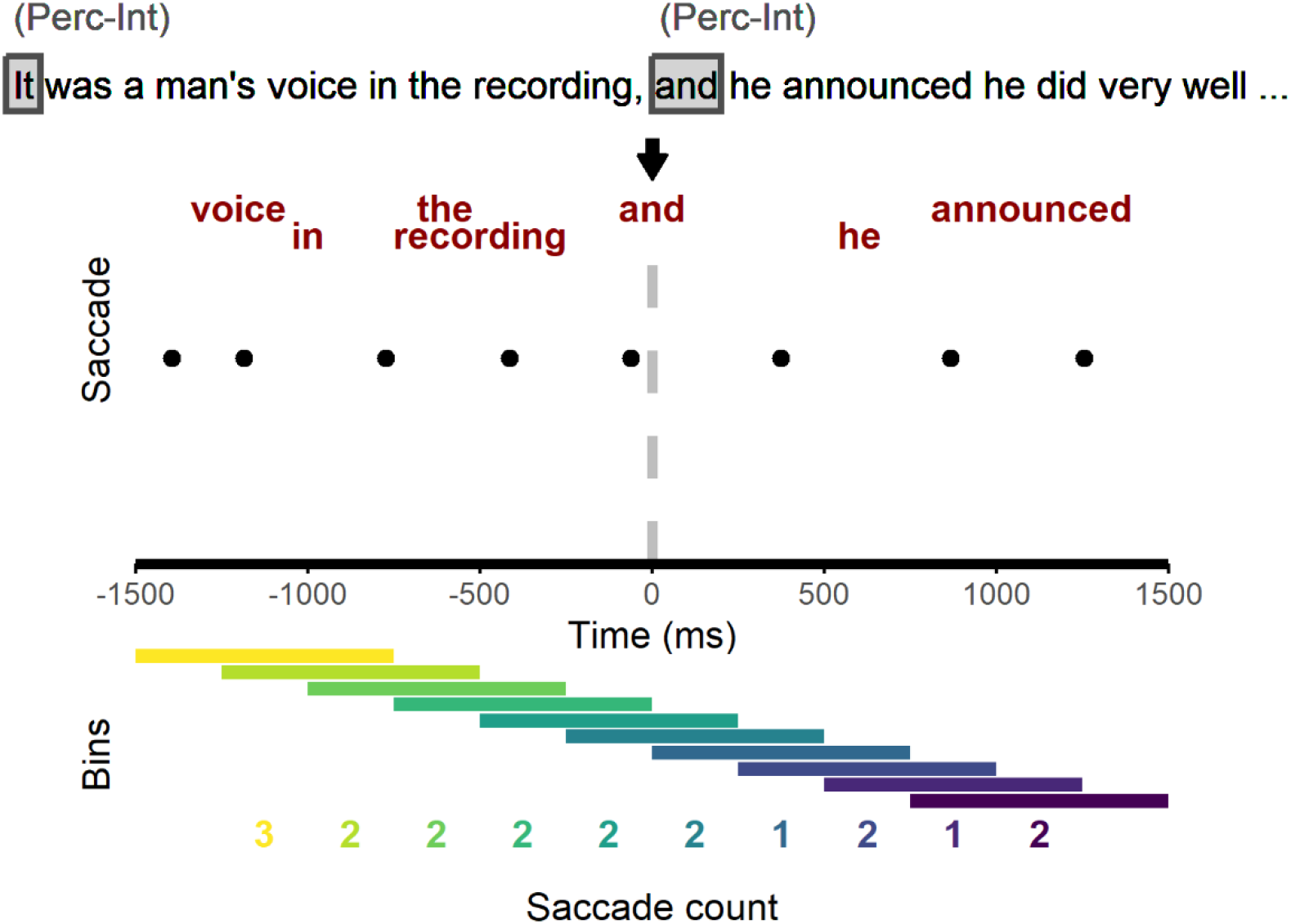
Illustration of Data Preprocessing into Sliding Time Windows *Note*. Text at the top of the figure represents a segment of the scored AI transcript, with an internal (perceptual) detail beginning at the word “It” and an internal (perceptual) detail beginning at the word “and.” The red text below demonstrates the timing of words surrounding the second detail within a 3000 ms time window. Dots below the words indicate saccade onsets. Saccades were summed in 10 time bins occupying the 3000 ms window. Each colored bar represents a time bin with the number of saccades appearing below in the same color, with bright/warm to dark/cool colors representing early to late time bins relative to the identified second detail.

