## Supplemental Information for "Remembrance with gazes passed: Eye movements precede continuous recall of episodic details of real-life events"

### Participants

Participants who reported significant medical or mental health complaints or prior exposure to Baycrest on initial online screening were excluded. As participants were originally recruited for a study on individual differences in episodic autobiographical memory (Armson et al, 2021), they were selected such that they reflected a full range of scores on the episodic subscale of the Survey of Autobiographical Memory (SAM; Palombo et al., 2013). This method of selection does not affect generalization of the present findings as the distribution of SAM episodic scores was normal, and the results were unaffected by inclusion of SAM-episodic scores in the model (see below).

### Procedure

The audio guide comprised 34 tracks that were initiated by the participant by clicking a button after the end of a preceding track (tour items are publicly available on the Open Science Framework https://osf.io/j25y4/). At each target item, participants were given the opportunity to examine the item and could also be directed to read an information card that provided information about the subject. The duration for examination of target items was controlled at 10 s. Participants were given extensive instructions before the tour began, and they were given an opportunity to practice using the MP3 player to control the audio guide. The tour was completed in two 10-minute segments, each occurring in a separate interconnected building. The segments were separated by a 45-minute battery of tests. Each section of the tour took approximately 10 minutes. Participants returned to the lab one week later whereupon they freely recalled the tour while eye movements were monitored under free- and fixed-viewing conditions, counterbalanced across the two tour segments. The current analysis only considered data during the free viewing condition.

Free recall responses were audio recorded and subsequently transcribed. Transcripts were scored using the Autobiographical Interview procedure (Levine et al., 2002), which categorizes details as internal (i.e., information directly relating to or contextually embedded in the event recollected) or external (i.e., semantic information or information not specific to the event recollected). Internal detail composites comprise event, place, time, perceptual, or thought details. External detail composites comprise semantic facts, repetitions, or other metacognitive statements or editorial comments. Five participants with fewer than 10 internal details were excluded as their recollections lacked sufficient richness to estimate a reliable relationship with eye movements. Scoring was performed by trained scorers who had achieved a criterion intra-class correlation of at least .90 on separate data.

### Eye tracking

Eye movements were recorded using an EyeLink II system (SR Research Ltd; Mississauga, Ontario, Canada). Eye tracking was accomplished via head-mounted video cameras that recorded positional data in the X and Y axis at a sampling rate of 500 Hz and spatial resolution of < 0.1 degrees. Head position was tracked by a single camera on the head mount that transmitted infrared markers to four sensors located on the corners of the computer monitor. Two additional camera mounts were located beneath each of the eyes and used infrared illuminators to measure eye positions via corneal and/or pupil reflections. The headband was padded and adjustable for participants' comfort, and the system could accommodate eyeglasses and contact lenses. Conversion of the eye movement data to a series of fixation and saccade events that were time-locked to memory recall (via an auditory tone) was achieved and interrogated with Data Viewer (SR Research Ltd.). Fixations and saccades were defined via EyeLink's online parser. If the velocity of two successive eye movement samples exceeded 30 degrees per second over a distance of 0.1°, the samples were labeled as a saccade. If the pupil was missing for 3 or more samples, the eye activity was marked as a blink within the data stream. Non-saccade and non-blink activity were considered fixations.

**Data preprocessing**

The free recall audio files were trimmed (removing instructions and prompts) and converted to a timestamped written transcription using Google Speech-to-Text in Python (https://cloud.google.com/speech-to-text), producing a transcript with the precise timing of each word at the millisecond level (500 Hz sampling rate). A custom MATLAB script merged the eye tracking and AM recall data on the same time series. Internal and external detail categorization from the separately transcribed and reliably scored protocol was then manually added to the time series. The AI manual defines a detail as a narrative element indicating a distinct idea or transition, which is typically at the beginning of a grammatical clause (a subject and a predicate; i.e., "… the sculpture was made of metal…") or a meaningful unit external to the clause (Seixas Lima et al., 2020). Accordingly, the boundaries of details were determined by conjunctions (e.g., "and," "but," or "so") or where typical punctuation (i.e., commas, semi-colon, or colons) would indicate transitions in ideas (see Figure 1 for example).

A sliding window was constructed around each detail onset (see Figure 3). Saccades were summed within 750 ms-wide bins, with each bin separated by a lag of 250 ms. This sliding window began at -1500 ms and continued up to +1500 ms, relative to the detail onset, for a total of 10 bins. As saccades persist over the course of milliseconds and therefore may occur within more than one time bin, a saccade contributed to a bin's sum if the timestamp associated with the saccade's initiation occurred within that bin.

**Analysis**

The relationship between eye movement timing and AM detail production was assessed using a logistic mixed-effects model (random intercept and slope) with the lme4 package (Bates et al., 2015) in R. Detail type (coded such that external details = 0 and internal details = 1) was regressed onto number of saccades and time bin. Our hypothesis was that saccade probability would increase prior to internal—but not external—detail generation; modeling detail type as a binary outcome thus allowed us to identify saccade patterns linked to episodic memory content over and above other verbal responses. Saccade counts were mean centered within each participant, so coefficient estimates reflected deflections from participants' average eye movements. Helmert contrast coding was used for time bin effects in all models. Regression coefficients represent the effect of each time bin compared to the average of all subsequent time bins. This contrast method was optimized to the sequentially ordered nature of the time bins to assess the temporal evolution of eye movements in relation to internal and external detail production.

Given the observed pattern of saccade amplification immediately before internal detail generation, exploratory mixed-effects (random intercept and slope) cubic polynomial regressions were conducted on internal and external detail data subsets to better characterize the nature of the distribution of saccades in relation to uttered details. Number of saccades was regressed on to linear, quadratic, and cubic terms for time bin.

**Individual differences: trait episodic autobiographical memory**

The present analyses were not concerned with individual differences in episodic autobiographical memory capacity, as was the case for the original study (Armson et al., 2021). Nonetheless, given that we previously found that SAM-episodic subscale scores moderated the relationship between saccades and internal details (Armson et al., 2021), we considered this factor in the logistic model as an ancillary analysis for completeness.

All effects previously reported by Armson et al, 2021 held when the relative timings of eye movements within the 3000 ms window were included in a linear mixed-effect model irrespective of time bins (random effects for intercept and saccade slope for participant), including the three way interaction whereby eye movements were related to internal (but not external) detail production in individuals with high trait episodic autobiographical memory as measured by the SAM-episodic subscale ($F(1, 1610.59) = 39.12$, $p < .001$). As in Armson et al, 2021, eye movements were significantly related to total detail production ($F(1, 64.05) = 4.93$, $p = .03$). Eye movements were also more strongly related to internal than to external detail production ($F(1, 1691.41) = 28.69$, $p < .001$), which was not evident in the original model, possibly due to the greater sensitivity of the present model in constraining variance to the 3000 ms window surrounding detail production.

When time bin was considered, SAM-episodic scores did not significantly moderate the relationship between early eye movements and internal detail recall (*p*s > .05), nor did the inclusion of the SAM-episodic subscale improve the fit to the cubic model ($AIC_{noSAM} = 104169$, $AIC_{SAM} = 104174$, $BIC_{noSAM} = 104236$, $BIC_{SAM} = 104276$; $\chi^2(4) = 2.40$, $p = .66$). We speculate that such individual differences operate at a coarser level of the retrieval epoch as opposed to the fine-grained level of the present analysis. In other words, individuals with low episodic AM capacity are less likely to retrieve internal details compared to those individuals with high episodic AM capacity (Petrican et al., 2020; Sheldon et al., 2016) but when they do retrieve such details, their eye movements follow the same temporal dynamics with respect to detail production.

Armson, M. J., Diamond, N. B., Levesque, L., Ryan, J. D., & Levine, B. Vividness of recollection is supported by eye movements in individuals with high, but not low trait autobiographical memory. *Cognition*, 206, 104487 (2021).

Bates, D., Maechler, M., Bolker, B., & Walker S. (2015). Fitting Linear Mixed-Effects Models Using lme4. *Journal of Statistical Software, 67*(1), 1-48. 10.18637/jss.v067.i01.

Palombo, D. J., Williams, L. J., Abdi, H., & Levine, B. (2013). The survey of autobiographical memory (SAM): a novel measure of trait mnemonics in everyday life. *Cortex, 49*(6), 1526–1540. 10.1016/j.cortex.2012.08.023.

Petrican, R., Palombo, D. J., Sheldon, S., & Levine, B. (2020). The Neural Dynamics of Individual Differences in Episodic Autobiographical Memory. *eNeuro, 7(*2), ENEURO.0531-19.2020. 10.1523/ENEURO.0531-19.2020.

Seixas Lima, B., Graham, N.L., Leonard, C., Levine, B., Black, S.E., Tang Wai, D.F., Freedman, M., & Rochon, E. (2020). Impaired coherence for semantic but not episodic autobiographical memory in semantic variant primary progressive aphasia. *Cortex*. *123,* 72-85. 10.1016/j.cortex.2019.10.008

Sheldon, S., Farb, N., Palombo, D. J., & Levine, B. (2016). Intrinsic medial temporal lobe connectivity relates to individual differences in episodic autobiographical remembering. *Cortex, 74,* 206–216. 10.1016/j.cortex.2015.11.005.
